# Genomic refugium of pre-domestication lineages in the Bronze Age Carpathian Basin

**DOI:** 10.1101/2023.06.29.547029

**Authors:** Zoltán Dicső, Géza Szabó, Róbert Bozi, Noémi Borbély, Botond Heltai, Gabriella Kulcsár, Balázs Gusztáv Mende, Viktória Kiss, Anna Szécsényi-Nagy, Dániel Gerber

## Abstract

Horse domestication is a key element in history for its impact on human mobility and warfare. There is clear evidence for horse control from the beginning of the 2^nd^ millennium BCE in the Carpathian Basin, when antler cheekpieces appear in the archaeological record mostly in the eastern areas. Previous archaeogenomic studies also revealed that the spread of the ancestors of modern day horses began at this time period, but the details of this event in Bronze Age Europe is yet to be uncovered. In this study we report a new shotgun genome (∼0.9x coverage) of a Middle Bronze Age horse (radiocarbon dated to 1740-1630 cal. BCE) from Tompa site, southern Hungary, along with six mitochondrial genomes from various sites from Late Copper Age to Early Bronze Age Western Hungary. Our results reveal a strong bottleneck among pre-domestication Carpathian Basin horses and delayed DOM2 introduction into the region compared to the surrounding areas. The population size reduction was most probably due to human mediated loss of natural habitat, but the practice of horsekeeping after the turn of the 2^nd^ millennium BCE can not be excluded based on the genomic data. Our results provide a complex history for horse domestication in the Central-European region, highlighting the need for further research to fully understand the extent and nature of human-horse interactions in this area throughout prehistory.

## Introduction

The horse is one of the most impactful and versatile animals ever domesticated, which enhanced workforce, warfare and human mobility. Still, our knowledge on the process and stages of horse domestication is incomplete. Archaeological evidence points to the ‘classical’ utilization of these animals at the beginning of animal husbandry through the findings of Botai (Kazakhstan) or Dereivka (Ukraine) sites, which suggest a highly horse oriented diet in the domesticator societies (Outram et al., 2021). However, the proposed success of herding, milking and their use as workforce (Gaunitz et al., 2018) were recently challenged by (Taylor & Barrón-Ortiz, 2021). Genetic evidence revealed that these horses were not the ancestors of modern livestock, and they represent a rather dead-end of domestication (Librado et al., 2021). Further indications for the presence and nature of utilization are sparse, ambiguous and indirect such as the anthropological analysis by (Trautmann et al., 2023) who have recently suggested that the practice of riding began in the 3^rd^ millennium BCE among Yamnaya culture associated societies. In the long and multi-stage process of domestication, there is clear archaeological evidence for carriage horses from the end of the 3^rd^ millennium BCE, whereas there is evidence for riding and milking from the first half of the 2^nd^ millennium BCE (Bozi & Szabó, 2023; Chechushkov et al., 2018, 2020; Maran, 2020; Wilkin et al., 2021).

In the Carpathian Basin, and especially in Transdanubia (Western Hungary) the archaeozoological data implies that during the Copper Age (∼3600-2800 BCE) in Boleráz and Baden cultures horse remains are rare, as e.g. at Balatonőszöd site, where only ∼0.6% of the recorded animal bones belonged to horses. In the following millennium the settlements of early Bronze Age Somogyvár–Vinkovci (∼2600-2200 BCE), Kisapostag/Encrusted Pottery cultures (∼2200-1500 BCE) (Gál, 2017) also yielded similar amounts of horse remains (between 0.5 and 1.43%, up to 5% in the Middle Bronze Age), with slight increase in frequency.

As opposed to the more scattered assemblages in other parts of Hungary, the first true evidence for horsekeeping in the region appeared on Bell Beaker culture (BBC) settlements around Budapest (e.g. Albertfalva site) towards the end of the 3^rd^ millennium BCE. Here the ratio of horse remains was very high (up to 60%), raising the possibility of a domestication center, but this theory is questioned by the large number of young animals that most likely kept for their meat (Dani & Kulcsár, 2021). The earliest direct evidence for horse herding through artifacts, such as antler cheek-pieces, linked to riding and/or other equine utilization appears in the archaeological record from the eastern part of Hungary from the 2^nd^ millennium BCE from the settlements of the Vatya and Füzesabony cultures (Bozi & Szabó, 2023) (Figure 1.).

**Fig. 1.**
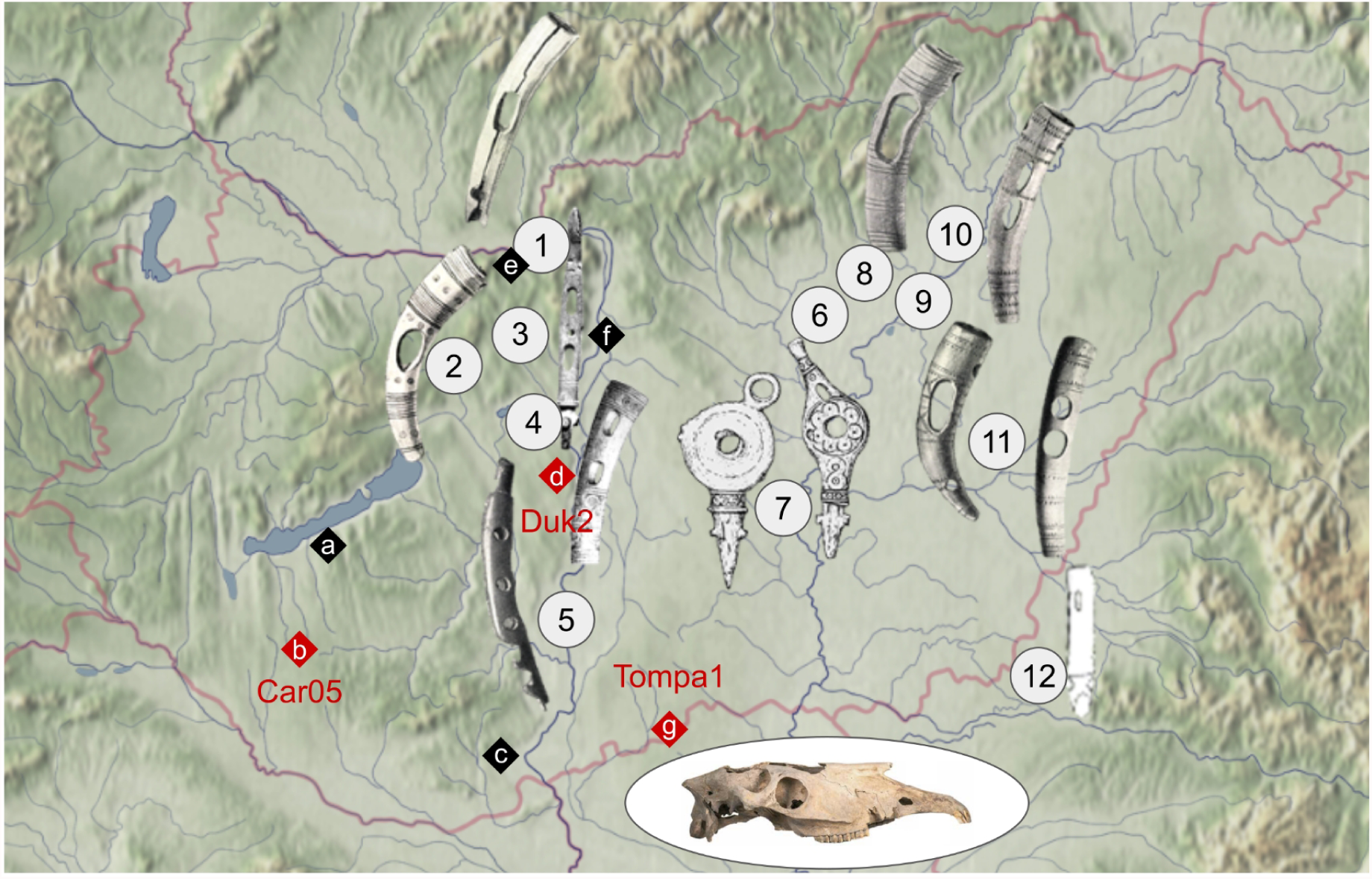
Sites in Hungary where horsekeeping related artifacts (antler cheek-pieces) were recovered from the Central European Middle Bronze Age (2^nd^ millennium BCE) 1 - Szob-Kálvária, 2 - Pákozd-Várhegy, 3 - Budapest-Lágymányos, 4 - Százhalombatta-Földvár, 5 - Gerjen, 6 - Jászdózsa-Kápolnahalom, 7 - Tószeg-Laposhalom, 8 - Füzesabony- Öregdomb, 9 - Tiszafüred, 10 - Mezőcsát-Pástidomb, 11 - Köröstarcsa, 12 - Pecica (Romania). Rhombus shaped marks denote the sampled sites (see Table 1 and Figure 2.), red rhombuses are for whole genome data: a - Ordacsehi-Bugaszeg, b - Kaposújlak- Várdomb, c - Szűr-Cserhát, d - Dunaújváros-Kosziderpadlás, e - Pilismarót-Basaharc, f - Budapest-Albertfalva, g - Tompa.

Earlier genomic studies that relied mainly on the hypervariable region of mitochondrial genomes were inept to reveal detailed population histories due to poor phylogeographic signals, although major trends, such as shifts in major genomic compositions were shown (Guimaraes et al., 2020). In recent years, whole genomes of ancient horses have revealed temporal and spatial distribution of dominant lineages, and the draft history of their domestication (Fages et al., 2019; Gaunitz et al., 2018; Librado et al., 2021). Accordingly, at least six major phylogenomic clusters existed in the past ∼100,000 years, from which only two exist today, the so-called DOM2, where all modern domestic breeds belong, and the descendants of Botai/Borly horses, known today as Przewalskii (Gaunitz et al., 2018), where the mongolian wild horses belong. Europe was dominated at least until the beginning of the European Bronze Age by two related and now extinct lineages, the Iberian and the Corded Ware culture (CWC). The CWC was named after the Corded Ware culture in (Librado et al., 2021), and we refer to this ancestry simply as Central European in this study. Horse remains with DOM2 ancestry are detectable from the Eneolithics in Yamnaya culture related archaeological context in Eastern Europe (Librado et al., 2021). These steppe groups’ westward expansion around ∼3000 BCE and their direct transmission into the CWC mediated a strong cultural and genetic impact on other European societies as well (Allentoft et al., 2015; Haak et al., 2015), but without the introduction of DOM2 horses until a thousand years later. Towards the end of the 3^rd^ millennium BCE ancient Central European horses began to be intensively replaced by DOM2, however, the details of this alternation is yet to be uncovered. Interestingly, the recently extinct tarpan horse has been revealed to be a 1:2 mixture of Central European and DOM2 groups, suggesting a more complex early population history for domesticated horses in Europe than other parts of the World (Librado et al., 2021). The Carpathian Basin in East-Central Europe is only represented in previous studies by two ancient horse samples (Car05, cal. 2571-2344 BCE; Duk2, cal. 2140-1977 BCE) from Hungary, both showing ancient Central European genomic makeup (Librado et al., 2021). In this study our aim is to unveil details on population structure and possible horsekeeping practices of local horses from the Late Copper Age to the Middle Bronze Age in the Carpathian Basin by involving new samples in the analyses.

## Results

In this study we present a newly sequenced ∼0.9x average coverage complete horse shotgun genome (Tompa1) from Tompa archaeological site from southern Hungary dated to 1740-1630 BCE, five newly sequenced and one resequenced (Car05) mitochondrial genome from Late Copper Age to Early Bronze Age (ca. 3600-1900 cal. BCE) from the area of today’s western Hungary (see Materials for site description, Table 1, Supplementary Table S1 and Figure 2.a).

**Fig. 2.**
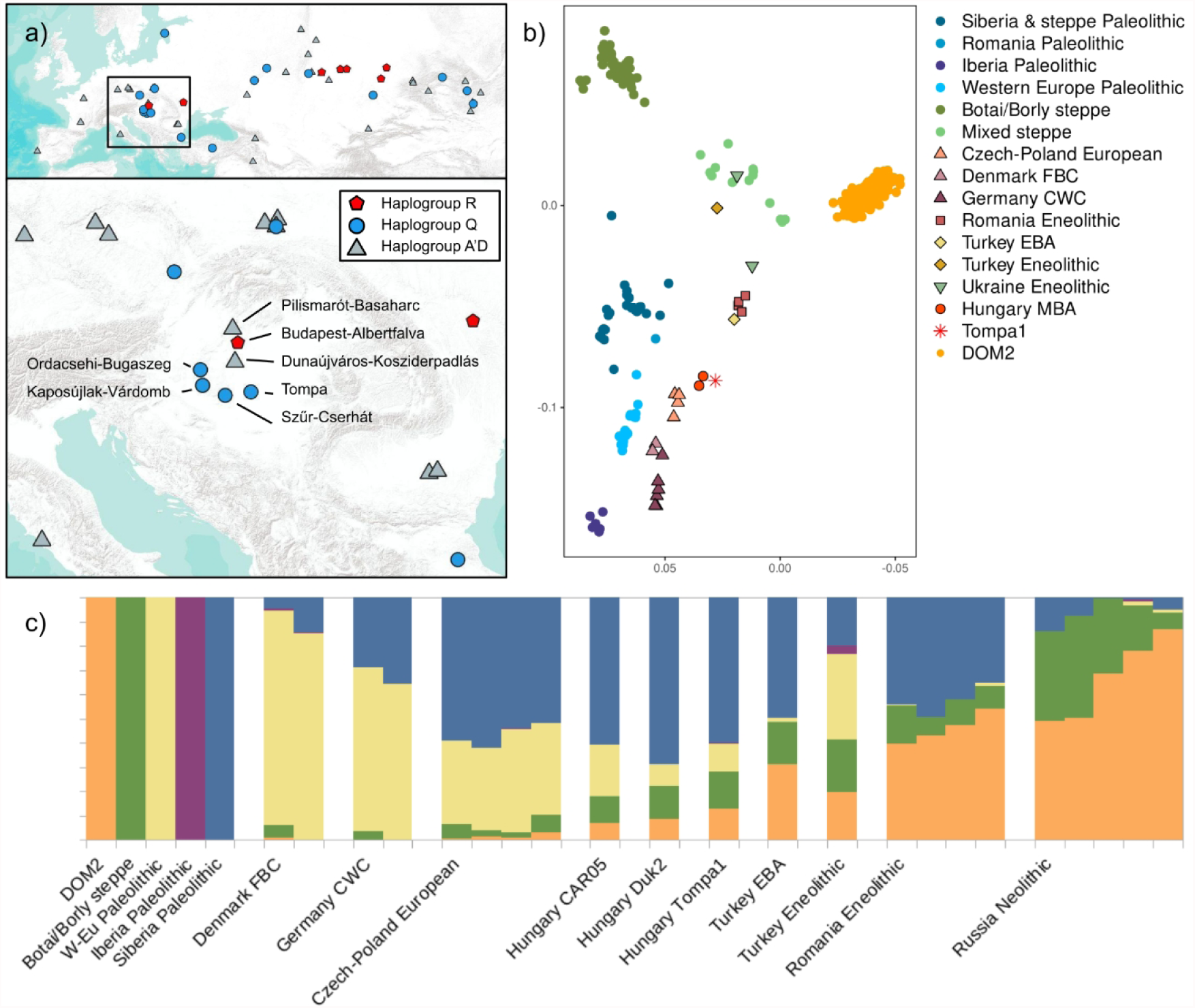
Genomic makeup of ancient horses from the Carpathian Basin a) Geographic distribution of mitochondrial haplogroups detected in this study b) PCA of ancient horse genotypes, prehistoric horses from Hungary have an intermediate location between Northern and Southern populations c) Struct-f4 analysis with 5 K positions Tompa1 between prehistoric horses from Romania and Hungary.

**Table 1.** Information on horse specimens presented in this study. mtHG means mtDNA haplogroup. Bold highlight denotes samples with whole genome data, and the ones with asterisk database data.

| Site (all in Hungary) | Culture | <sup>14</sup>C/cal BCE (1σ) | Sample ID | mtHG |
| --- | --- | --- | --- | --- |
| Pilismarót-Basaharc | Baden | 3630-3530 | Car03 | A'D |
| Szűr-Cserhát | Baden/Somogyvár-Vinkovci | 2880-2680 | Car04 | Q |
| **Kaposújlak-Várdomb** | **Somogyvár-Vinkovci** | **2570-2460** | **Car05\*** | **Q** |
| Kaposújlak-Várdomb | Somogyvár-Vinkovci | 2560-2410 | Car06 | Q |
| Budapest, Albertfalva-Hunyadi J. u. | Bell Beaker | 2280-2150 | Car07 | R |
| **Dunaújváros-Kosziderpadlás** | **Nagyrév** | **2140-1970** | **Duk2\*** | **A'D** |
| Ordacsehi-Bugaszeg | Kisapostag | 2010-1910 | Car08 | Q |
| **Tompa** | **Vatya** | **1740-1630** | **Tompa1** | **Q** |

### mtDNA composition

We created a mitochondrial haplogroup classification tool freely available at https://github.com/ArchGenIn/cablogrep called *cablogrep*, with a similar workflow to the popular tool Haplogrep (Weissensteiner et al., 2016) developed for human mtDNA. The adapted haplogroup system is based on (Achilli et al., 2012) supplemented by haplogroups X and S from (Der Sarkissian et al., 2015) and (Guimaraes et al., 2020). We restricted the mtDNA analysis to sequences that have less than 50% missing data (N or ambiguous base). Contrary to the modern day horses, maternal lineages of prehistoric livestock and wildlife turned out to be informative from a phylogeographic perspective in East-Central Europe. Most of our samples can be divided into two major haplogroup clusters: Car03 and Duk2 belong to mtHG A’D and the others to mtHG Q, except Car07, which has mtHG R (Table 1, Figure 2.a). This composition is strikingly homogeneous compared to populations from the surrounding areas (Guimaraes et al., 2020; Librado et al., 2021).

Geographical and chronological distribution of both mitochondrial haplogroups Q and A’D seems to be structured, as per our data haplogroup Q concentrates to today’s Hungary and can be assumed in prehistoric Southeast Europe too, as it appeared mostly in Anatolian horses after 2000 BCE (Guimaraes et al., 2020). On the other hand, haplogroup A’D was more frequent outside of the Carpathian Basin, although both were found equally across the Pontic-Caspian steppes. Haplogroup R, however, is almost ubiquitous to Botai horses, but also appeared in Bronze Age Volga-Don area, making its appearance inside the Carpathian Basin noteworthy and the connection of Car07 to steppe horses probable, especially in the light of a contemporaneous DOM2 horse in today’s Czechia (2200–2000 BC, Únětice culture from Holubice site (Librado et al., 2021)).

### Genomic makeup of Tompa1

A set of ∼1,6 million biallelic SNPs were considered for population genomic analyses, obtained from (Jagannathan et al., 2019), after applying further filters (for details, see Methods). Principal component analysis (PCA, Figure 2.b) reveals major clines among ancient and extant horses. Tompa1 clusters with the two other Hungarian Bronze Age horses, situated in an intermediate position between Neolithic and Eneolithic Turkish/Romanian and Czech/Polish horses, reflecting the natural genomic cline of wild horses in the region (Librado et al., 2021). *Struct-f4* analysis (Librado & Orlando, 2022) with 5 components also confirms the position of Tompa1 (Figure 2.c), although it shows some similarity to Eneolithic Romanian and Turkish samples through an elevated component modeled by DOM2 horses. *f*3 analysis (Maier et al., 2021; Patterson et al., 2012) shows highest similarity of Tompa1 to Duk2 and Car05 (merged into one Hun_MBA group, Supplementary Table S2), but *f*4 statistics (Maier et al., 2021; Patterson et al., 2012) in form of *f*4(Tompa1, Hun_MBA, DOM2, Donkey) revealed further affinity towards steppe horses. To assess whether this extra DOM2-like component is the sign of actual mixture between local and steppe stocks or the sign of the geographic cline we applied qpAdm (Haak et al., 2015; Maier et al., 2021; Petr et al., 2019) analysis. Tompa1 can be modeled with qpAdm as two-way mixture of Hun_MBA (c. 95-78%) and an Eastern component, providing the highest estimates for Turkey_EBA (22.1±7.84%, *p*=0.154) and only 4.78±3.06%, *p*=0.272 for DOM2 as fitted sources. While three-way admixture models of Hun_MBA, Turkey_EBA and any other steppe related population yield acceptable *p*-values, the third components are too small and standard errors are too high to consider it reliable. These results are on par with Turkey_EBA second position on *f*3 and previous results on Hun_MBA (Librado et al., 2021), except Tompa1 has slightly higher levels from this Anatolian-related ancestry. Considering that Tompa1 was found on the lowest longitude among Carpathian Basin horses, all these results robustly position it on a natural genetic cline between Anatolia and the Carpathian Basin.

Considering the occurrence of haplogroup R in Bell Beaker cultural context in northern part of Central Hungary (around Budapest) and whole genome evidence for DOM2 introduction in Czechia ∼300-400 years prior Tompa1, as well as the possible modeling of Tompa1 with minor DOM2 ancestry, we evaluated the origin of this component further, in order to determine whether it is part of the wild genetic structure of horses in the area, i.e. part of the NEO-ANA cline described in (Librado et al., 2021), or whether it is the sign of minor but recent introduction of DOM2 ancestry to an already existing local livestock.

In order to test this, we applied *Admixfrog* (Peter, 2020) software to assess DOM2- like genomic chunk sizes in Tompa1 and Hun_MBA horses (see Supplementary Table S3), as well as for sample Kan22 (Turkey_EBA) for comparison (for details, see Methods). Accordingly, Hun_MBA and Tompa1 horses do not show particular ‘aggregation’ of steppe related genomic chunks; instead, steppe related chunks are distributed evenly in small sizes roughly equally in homozygous and heterozygous form, in complete par what one would expect from a geographic-cline induced, over longer time manifesting genetic drift (Figure 3).

**Fig. 3.**
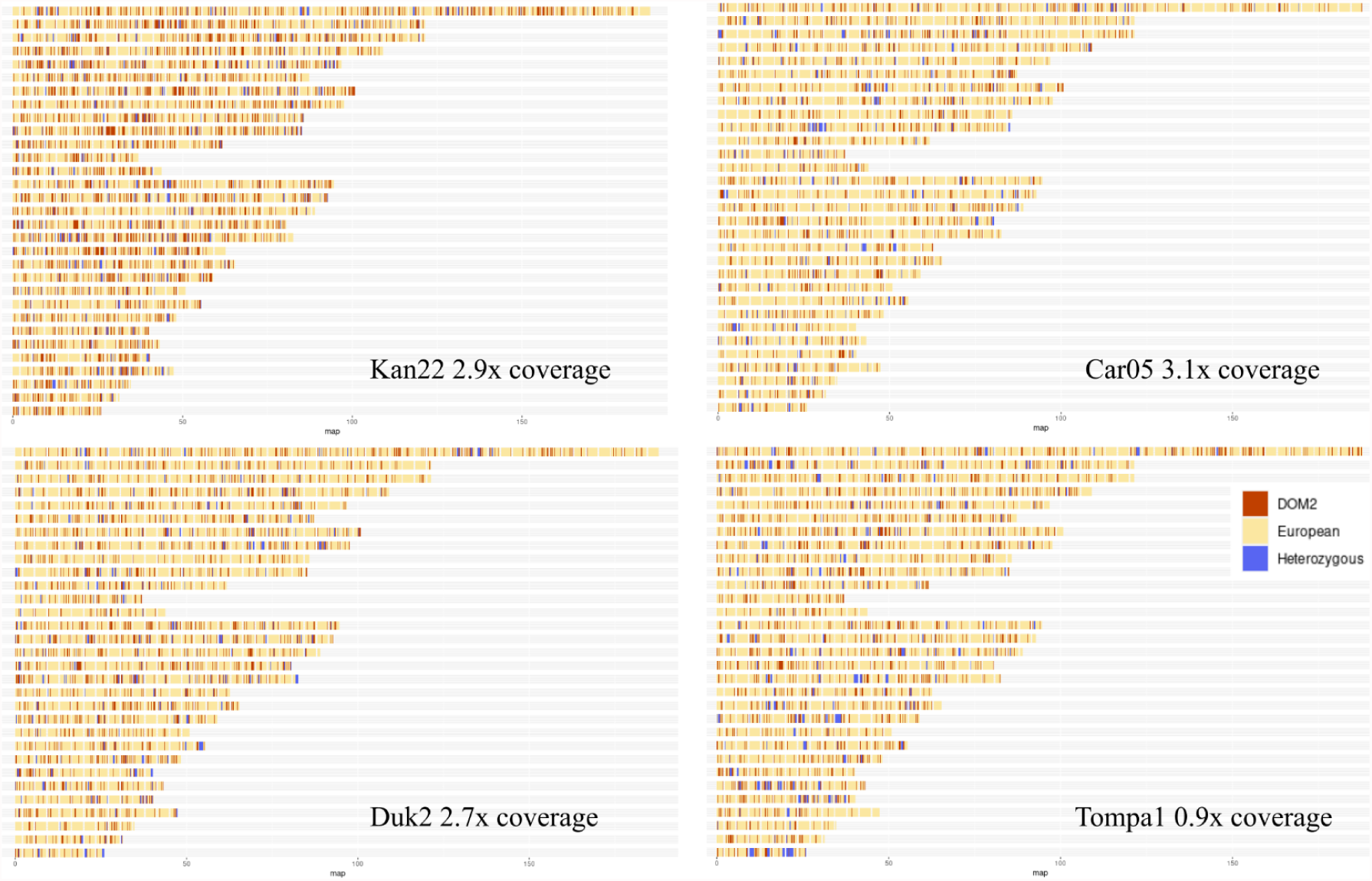
DOM2 genomic segments of the investigated horses, analyzed with Admixfrog (Peter, 2020). Kan22 (Turkey EBA), Car05, Duk2 and Tompa1 (Hun_MBA) are equally shown highly fragmented DOM2-like ancestry, suggesting no recent DOM2 introduction to these populations.

### Genetic traits

There are a number of genetic traits that are routinely screened in horse pedigree analyses, mostly related to physical appearance, temper and inheritable diseases (Fages et al., 2019; *Online Mendelian Inheritance in Animals, OMIA. Sydney School of Veterinary Science*, n.d.; Srikanth et al., 2019). We made an arbitrary selection of frequently measured variants (variant list and results are in Supplementary Tables S4a-c) that we tested on Hun_MBA and Tompa1 with a custom script, using a variant detection threshold system described in (Gerber et al., 2022). Accordingly, these horses mostly possessed the ancestral variants for traits related to temper, although variants on *ACN9*, *CKM*, *COX4/1* and *COX4/2* genes that are associated with racing performance show mostly derived variants in all three horses from Hungary (if the loci were covered), in line with observations and most probably the interpretation of (Schubert et al., 2014) of these being present in European horse populations since the Paleolithic. Variants on the *ZFAT* gene, however, show a completely opposite set in Car05 compared to Duk2 and probably to Tompa1 (only 1 variant is covered for this specimen), raising the possibility of selection against smaller wither height between ∼2500-2000 BCE. Interestingly, Car05 is also heterozygous to the silver coat coloration, and despite the base color could not have been identified as no reads covered the *ASIP* gene, it can be assumed that specimens with silver coat were present in this horse population.

Regarding traits associated with inheritable diseases, Duk2 carries the malignant variant for recessively inherited chondrodysplasia (*SLC26A2*), characterized by abnormal body proportions and dwarfism (Hansen et al., 2007). The variant has a coverage of 3x, all with the malignant substitution, and it is positioned around the border of a presumed ROH segment (see next section), thus it is fairly reasonable to suspect the actual onset of the symptoms.

We also performed *RAiSD* (Alachiotis & Pavlidis, 2018) analysis to search for signs of selective sweep among non-DOM2 horses around Europe. In this analysis we used Paleolithic to Bronze Age data from modern-day France, Eneolithic samples from Germany, Czech Republic, Poland, Hungary, Romania and Turkey. For this particular task we used a diploid dataset, which was generated with *bcftools* (Li, 2011) (for further details, see Methods). Our results revealed signals of selection among a *HOXB* gene cluster in chromosome 11 and the *PDE10A* gene in chromosome 31 (Figure 4). The former is associated with physical development and vertebral modifications, and is the same cluster that was highlighted by Fages et al., but we found no selection within the *HOXC* gene cluster. The *PDE10A* gene is associated with a variety of traits, and among others it is associated with control of movement and cognition (Diggle et al., 2016; Mencacci et al., 2016).

**Fig. 4.**
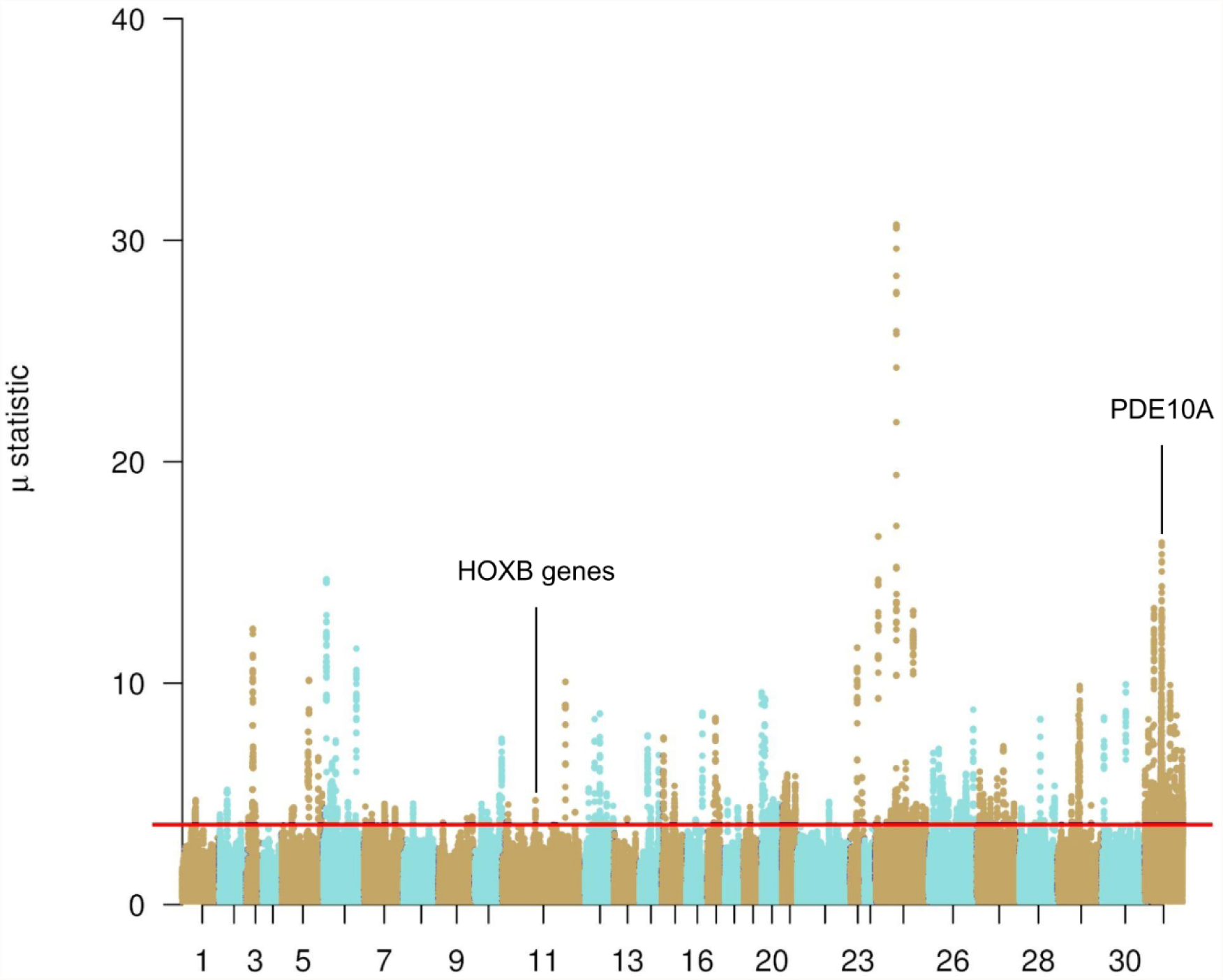
RAiSD (Alachiotis & Pavlidis, 2018) results show that HOXB gene cluster and PDE10A genes underwent selection among non-DOM2 horses in Europe.

### Population size

We performed runs of homozygosity (ROH) analysis by ROHan (Renaud et al., 2019) to assess the general pattern of inbreeding. Unfortunately, Tompa1 does not have enough coverage for proper analysis, but Car05 and Duk2 does, along with a number of ancient database samples. Since results tend to be higher ROH-skewed towards lower coverages, besides inferences on the original dataset and a set of relevant database samples (see Supplementary Table S5 for samples and results), we performed this test on subsampled (down to the coverage of Duk2, ∼2.7x) BAM alignment files as well, in order to see whether the general patterns across samples remain with lower coverage. Accordingly, a general pattern of increasing ROH percentages in time forward can be observed as a sign for a bottleneck that has already been shown by previous studies (Fages et al., 2019; Gaunitz et al., 2018; Librado et al., 2021). Surprisingly, Hun_MBA and Kan22 (Turkey_EBA) show the highest proportions for ROH segments (up to 8%), suggesting either extreme population size reduction or intensive selective breeding. While these extreme values also could be the result of lower coverage, the downsampled dataset does not show a significantly different distribution of ROH across the samples, although standard errors reach such a level that make individual evaluations ambiguous.

We supplemented this analysis with *ASCEND* (Tournebize et al., 2022), which aims to reveal the timing of founder-effects and genetic bottlenecks. While *ASCEND* is supposed to work properly on low coverage samples, it requires at least five individuals from a narrow chronological and geographical segment. This issue currently can not be overcome, but a simple experiment would reveal uneven timing of bottlenecks and general genetic diversity in the studied region: we merged relatively contemporaneous non-DOM2 horses from Hungary (Hun_MBA and Tompa1), Romania, Germany, Turkey (Kan22), Poland and Czechia, assuming that they belong to the general East-Central European horse population. We ran *ASCEND* with all samples - once with, once without Romanian horses, as these are much older than the others - then we excluded populations in each subsequent run. The pattern observed in ROH analysis is not entirely reflected by the *ASCEND* outputs (Supplementary Table S6). With Romania included (Figure 5.a) the exclusion of Hun_MBA shifts the bottleneck timing farther, suggesting that Hun_MBA added lower overall diversity to the group, suggestive for a separate bottleneck for this population compared to others in line with ROH observations, and the same applies to Germany CWC too, although for the high standard errors the results are rather tentative. Similarly, the exclusion of Romania Eneolithic shifts the bottleneck timing closer to the test groups, suggesting that their population is more diverse, which is suspected for their oldest chronological age among all individuals in this test. When we exclude Romania Eneolithic from the populations (Figure 5.b), the exclusion of Germany CWC but not the Hun_MBA changes the output significantly, suggesting an independent and more recent bottleneck event for the horse population in today’s Germany in the first half 3rd millennium BCE, and similarly dated but much severe bottleneck for Hun_MBA compared to other European populations.

**Fig. 5.**
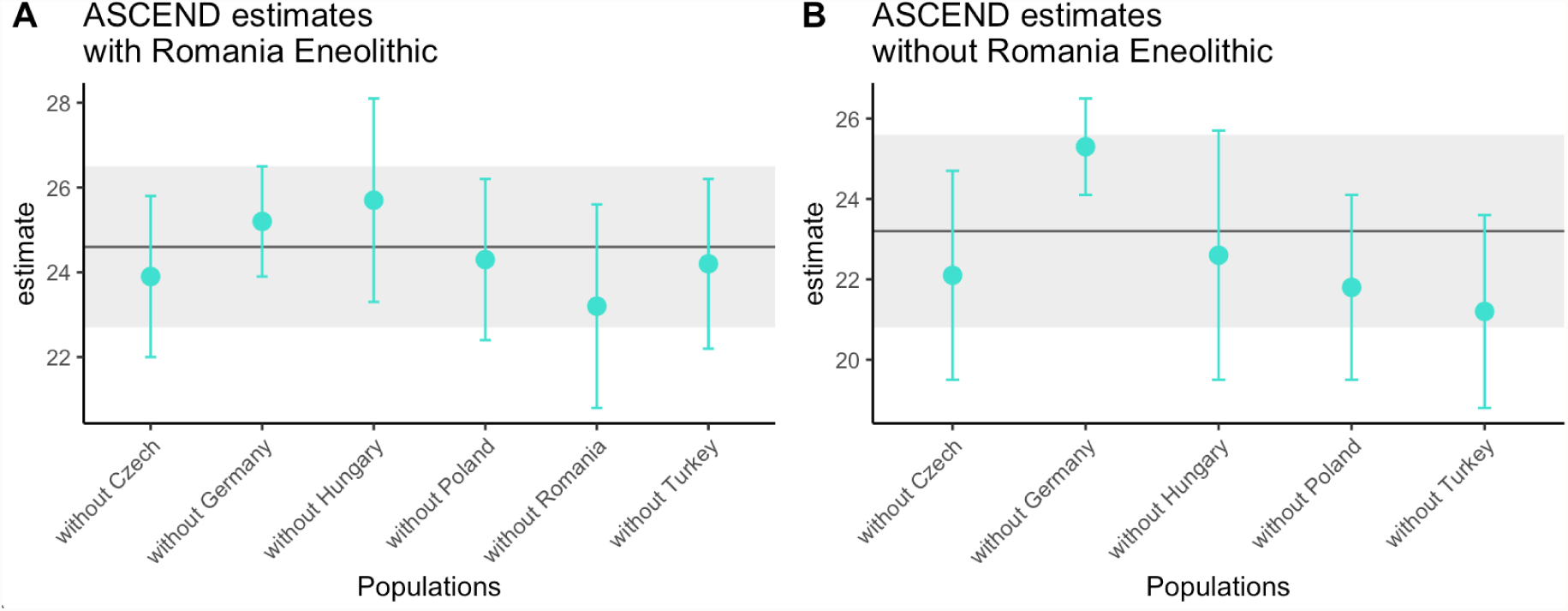
ASCEND (Tournebize et al., 2022) results with time estimated for founder-effects and genetic bottlenecks in samples that are relatively contemporaneous (dated between ∼2800-1800 BCE). The solid line represents the estimate for all populations and the gray bar its standard deviation. Data are shown in Table S6.

## Discussion

According to previous studies on Late Neolithic, Copper Age and Early Bronze Age Central Europe (Szabó, 2017), the region faced extreme changes connected to human mobility (Allentoft et al., 2015; Haak et al., 2015) and activities connected to agricultural advancements (Gamarra et al., 2018; Gulyás & Sümegi, 2011; Magyari et al., 2012; Reed, 2017). However, archaeogenomic studies on livestock and wildlife especially from these periods are very limited, and our knowledge on this topic is rather sparse. The mtDNA composition of prehistoric horses in the Carpathian Basin suggests a rather homogeneous population in the area during the studied period, which, or more likely its Southeastern European counterpart, may have contributed to at least some extent to the Anatolian livestock after 2000 BCE, as suggested by (Guimaraes et al., 2020). It is possible that DOM2 horses from the steppes were bred with local mares there, as mtDNA haplogroup Q marks this Anatolian replacement to a high extent, contrary to the general steppe composition of haplogroup A’D predominance over Q. However, based on the results of (Librado et al., 2021), the genomic contribution of these Southeast European livestocks to the Anatolian gene pool - at least for the autosomally tested specimens - must have been minor. On the other hand, in whole genomes of the Carpathian Basin the DOM2 contribution is not detected before 1740-1630 BCE in the region, pointing to the delayed replacement of the local population. This stands in contrast with the introduction of DOM2 horses to surrounding regions (today’s Czechia, northern part of Central Hungary, Moldavia and western Turkey) between 2200-1900 BCE in various cultural contexts, suggesting some sort of regional cultural or ecological isolation of yet unknown nature.

A number of archaeological (Dani & Kulcsár, 2021; Grigoriev, 2021; Kanne, 2022; Makarowicz et al., 2023; Przybyła, 2020), anthropological (Kanne, 2022; Trautmann et al., 2023), archaeozoological (Bökönyi, 1959, 1974, 1978; Choyke & Bartosiewicz, 2000, 2005; Vörös, 1988; Vretemark & Sten, 2005) and genetic (Librado et al., 2021) evidence point to at least partial domestication of non-DOM2 horses in Europe, including the studied area, but the extent and the start of this practice is highly disputable. As suggested by the archaeological evidence, the BBC could have had at least partially domesticated livestock in northern part of Central Hungary at the end of the 3^rd^ millennium BCE, and the mitochondrial haplogroup of Car07 horse specimen, that implies its (at least maternal) connection to DOM2, strengthens this hypothesis. On the other hand, results regarding the local non-DOM2 livestock remained slightly controversial. Severe bottleneck as indicated by ROH analysis as well as high homogeneity of the mitochondrial gene pool suggest highly decreased population size for horses in the Carpathian Basin and probably in the Balkans and Anatolia as well at the study period, but details of this process remains unclear. The possible selection for *HOXB* gene cluster, *PDE10A* and *ZFAT* genes propose a human mediated bottleneck at least towards the end of the 3^rd^ millennium BCE. However, 1) the fine-scale correlation between geographic location and genomic pattern of all samples, 2) non-significant differences in population size reduction between European populations (except for Germany CWC) in *ASCEND* analysis and 3) the lack of recent (human-mediated) DOM2 introduction all indicate a small but wild population in the region. The outstandingly high drop in the genomic diversity among CWC horses in Germany in the early 3^rd^ millennium BCE as a signal for their domestication is on par with them being the best proxy for the Central European component of the tarpan horse (Librado et al., 2021), and in our view the results are conclusive to assume actual domestication of that population, however, low data quality inhibits further tests for horse selection or inbreeding in the CWC context.

According to the available archaeological evidence (Bozi & Szabó, 2023; Dani & Kulcsár, 2021; Gál, 2017; Kanne, 2022), widespread practices for horsekeeping can not be observed in the local (Carpathian Basin) archaeological cultures until the end of the 3^rd^ millennium BCE. In our view, the similarly dated but much severe population size reduction in the Carpathian Basin (and possibly in the Balkans and Anatolia) compared to other regions of Europe can be traced back to several ecological factors and human mediated processes, but not to direct population control, such as domestication or overhunting. We suggest two major hypotheses to explain the results:

1. Towards the end of the Ice Age, the reforestation of Europe occurred much later in the North European Plain than in Southeast Europe and the Carpathian Basin(Birks & Willis, 2008; Roberts et al., 2018; Svenning et al., 2008), thus habitat fragmentation in the study area likely caused a natural bottleneck for horses. This could have resulted in a metapopulation structure of scattered smaller groups with higher inbreeding levels, that could explain both ASCEND and ROH results. Accordingly, no particular human-mediated effects are required to explain the different bottleneck in the Carpathian Basin, as the same Neolithisation process across Europe met an already decreased population in the study region.
2. Based on several studies (Baldia et al., 2013; Betti et al., 2020; Müller, 2013; Porčić et al., 2016, 2021; Shennan et al., 2013; Timpson et al., 2014), the region witnessed a generally more extensive human settlement activity than other parts of Europe during the Neolithic, which could have affected local horse populations more drastically via loss of habitat and maybe hunting (Gál, 2017).

Certainly, these processes could jointly affect the changes in horse populations, but further data is required to verify these hypotheses. Nevertheless, our data do not evidently disclose small scale and sporadic local attempts for taming or utilization or the presence of some sort of selective breeding, but widespread herding practices similar to DOM2 horses in our view are highly unlikely (Bozi & Szabó, 2023; Gál, 2017). Further data and sampling is required for more firm evaluations, nonetheless our results contribute to the general understanding of horse herding practices in the Central-European region in the Bronze Age.

## Materials and Methods

### Materials and archaeological background

At Pilismarót-Basaharc a settlement and cemetery for the Baden culture was recovered. In grave 434 two articulated tarsal bones from the right limb of a horse were recovered, likely as an offering to the deceased. Based on the calcaneal tuber ossification, the animal was at least three years old. In the settlement remains of other animals, such as goats, dogs, deers, wild boars, etc. were also recovered (Gál, 2015).

At Szűr-Cserhát site a settlements of the Baden and the subsequent Somogyvár-Vinkovci cultures were recovered. Both periods yielded large amounts of cattle, goats and sheep remains, as well as some horse bone fragments. At object 37 a red deer skeleton was buried on top of the pit along with bones of cattle, caprines, pig, dog, wild boar and a single horse first phalanx (Gál, 2017).

At Kaposújlak-Várdomb settlements of several periods were excavated(Somogyi, 2004). There were no horse bones in the objects of the Copper Age archaeological horizon, but a total of 39 horse bones (0.8% of all animal remains) were identified among the Somogyvár-Vinkovci culture related findings. Most of these belonged to mature animals, but juveniles were also recovered (Gál, 2017).

At Budapest-Albertfalva site a settlement of a local variant (Csepel group) of BBC was recovered, where horse remains were present in such high proportions (41.83%), that is unparalleled at any contemporaneous sites in Central Europe at this period. The population of the village may have been at 50-60 people, there is an estimate of between 1.18 and 1.42 horses per person. The relatively large number of juveniles (∼31% of all horse remains) at Albertfalva has been suggested to indicate meat production. The bones were heavily calcined and fragmented for marrow extraction (Lyublyanovics, 2016).

At Ordacsehi-Bugaszeg site settlements of several periods were excavated. Horses were represented in ∼0.6% of the animal remains at the site; the sampled bone fragment from Feature 1470/2018 was among the mixed findings of Somogyvár-Vinkovci and Kisapostag cultures. Later, radiocarbon dating placed the bone fragment within the Kisapostag horizon. The majority of horse bones originate from fully grown specimens. Based on the epiphysis ossification level of the distal fragment of the femur the animal was younger than 4 years (Gál, 2017).

At Tompa archaeological site, which belongs to a settlement of the Vatya culture, a complete skull of a horse was recovered that belonged to an 8 years old mare, dated between 1740-1630 BCE. Based on the morphological analysis of (Bozi & Szabó, 2023), the specimen may have shown traces for nose bit wearing.

### Ancient DNA laboratory work

Laboratory work was performed under sterile conditions in a dedicated ancient DNA laboratory facility (Institute of Archaeogenomics, Research Centre for the Humanities, Eötvös Loránd Research Network, Budapest, Hungary). The laboratory work was carried out wearing protective clothing. Separated work areas were irradiated with UV-C light, surfaces were cleaned with DNA-ExitusPlus™ (AppliChem) and/or bleach.

Petrous bone and tooth were taken from Tompa1. Sample surfaces were cleaned by sandblasting and UV-C irradiations. Bone powder was used to extract DNA from the petrous bone, and from the cementum layers of the tooth. Two DNA extraction was carried out from each sample, blank controls were included at all steps. DNA extraction was performed according to (Rohland et al., 2018) on liquid-handling systems. DNA libraries were prepared using UDG-half treatment based on the protocol of (Rohland et al., 2015) with minor changes suited for automation on liquid-handling platforms. Unique double internal adapter combinations were used for every library. Libraries were then amplified with TwistAmp Basic (Twist DX Ltd) and purified with AMPure XP beads (Agilent). Libraries for shotgun sequencing were indexed using unique iP5 and iP7 indices (Meyer & Kircher, 2010). DNA concentration of each library was measured on Nanodrop (Thermo Scientific), fragment sizes were checked on Agilent 4200 TapeStation System (Agilent). NGS sequencing was done on an Illumina MiSeq System using the Illumina MiSeq Reagent Kit v3 (150-cycles) and on an Illumina Novaseq 6000 System (Novogene Company, China).

For mitochondrial DNA examination each cleaned sample (Car03-Car08) was ground down. DNA extraction was based on the protocol from (Dabney et al., 2013) using 110 mg of bone or tooth powder. Double-stranded libraries were prepared following the protocol of (Rohland et al., 2015), using internal unique barcodes. Libraries were then amplified with TwistAmp Basic (Twist DX Ltd) and purified with AMPure XP beads (Agilent). Mitochondrial RNA baits for DNA capture were prepared in-house following the protocol (SM2) by (Llamas et al., 2016) with modification for horses: present-day mitochondrial genome was amplified for baits in four overlapping fragments using the Expand Long Range dNTPack kit (Roche) with four selected primers pairs (175F-5151R, 4842F-8437R, 7978F-12894R, 12546F-563R) from (Achilli et al., 2012) with an annealing temperature of 60°C. For capture hybridization assay 300-500 ng of barcoded DNA library and 440 ng of biotinylated RNA baits were used. After the last wash step of the hybridization, beads were directly used for off-bead PCR amplification using the Phusion High-Fidelity PCR kit (New England Biolabs) for 30 cycles with primers from library amplification (Rd1, Rd2) in total volume of 25 µl. The DNA concentration of each captured library was measured on a Qubit 2.0 fluorometer (Life Technologies). Captured samples for shotgun sequencing were indexed using universal iP5 and unique iP7 indices (Meyer & Kircher, 2010) and purified with AMPure XP beads (Agilent). The libraries were then pooled in equimolar ratios. Sequencing was performed on an Illumina MiSeq System using the Illumina MiSeq Reagent Kit v3 (150-cycles) according to the manufacturer’s instructions.

### Bioinformatic analyses

#### Sequencing data processing

Raw sequencing reads were processed through the PAPline (Gerber et al., 2022) bioinformatic package with default options, where all of the sequencing libraries for a single sample were merged together. EquCab3 genome assembly (Kalbfleisch et al., 2018) was used as a reference for whole genome analysis; mitochondrial genomes were obtained through mapping reads to only the mitochondrial reference sequence from EquCab3. Species and genetic sex designation was performed with Zonkey (Schubert et al., 2017). We created a bioinformatic tool for mtHG classification for horses based on the system of previous studies (Achilli et al., 2012; Der Sarkissian et al., 2015; Guimaraes et al., 2020), for the detailed description of the software see github https://github.com/ArchGenIn/cablogrep.

#### Whole genome analyses

As a background for our analyses, we downloaded all publicly available ancient horse genomes (Fages et al., 2019; Gaunitz et al., 2018; Librado et al., 2021). After producing BAMs aligned to EquCab3.0 with *PAPline* (Gerber et al., 2022), we used *samtools* v1.10 (Li et al., 2009) to generate a pileup file. We obtained a list of variant positions available on the European Nucleotide Archive under project name PRJEB28306 (Jagannathan et al., 2019). This list contains ∼23 million genomic variants of 88, mostly modern domesticated horses, from which we selected ∼1.65 million representative variants by applying additional filters suitable for ancient DNA analysis. We kept only biallelic transversions of single nucleotide polymorphisms (SNP), and we further discarded positions under MAF 0.05, and positions with missing genotypes, and we set a minimum distance of 1000 bp between SNPs. Variants from the X chromosome were also excluded. We used this list of positions to generate the Eigenstrat (Patterson et al., 2017) format pseudohaploid genomes for population genomic analyses with *samtools* and *pileupCaller* v1.5.2 (Schiffels, 2018). For further necessary format conversions, we also used the *Eigensoft* (Patterson et al., 2017) software package and *Plink1.9* (Purcell et al., 2007).

*Smartpca* v16000 (Patterson et al., 2017) was used to perform PCA analysis, with shrinkmode and lsqproject option. The populations we used for smartpca: ancient DOM2, Russian Botai, Central steppe EBA, Hun_MBA, Iberian Paleolithic, France Paleolithic, Germany CWC, Ural Paleolithic, Poland FBC, Russia Paleolithic (*Equus lenensis*), Romania Eneolithic, Turkey EBA, Kazakh Tersek Eneolithic, Kazakh Borly, Czech Neolithic, Russia Taymyr Paleolithic, Russia Yamnaya. We used *Struct-f4* (Librado & Orlando, 2022) to infer ancestral proportions with K=5 for relevant populations, where we excluded samples with lower than ∼1x coverage. For *f*-statistics, we used *admixtools* (Maier et al., 2021; Patterson et al., 2012).

For *RAiSD* we used a diploid dataset. This dataset was generated with *bcftools.* We ran *RAiSD* with the following parameters: -M 3 -y 2 -A 0.995. The A parameter was the threshold for the Manhattan plot, the 0.995 value was the recommendation of the author.

## Acknowledgement

This paper was supported by the Momentum Mobility research project (LP 2015-3) hosted by the Institute of Archaeology, Research Centre for the Humanities, granted by the Hungarian Academy of Sciences. We would like to greatly thank archaeozoologist Erika Gál and archaeologist Mária Bondár for their assistance in providing samples and ^14^C dates to this paper.

## Author contribution

D.G. and A.Sz-N. conceived and designed the experiments. Z.D. processed the sequencing data and performed the analyses. D.G. and N.B. collected the mtDNA data and designed *cablogrep*. G.Sz. and R.B. provided the Tompa1 sample and jointly evaluated the archaeological context with G.K. and V.K. M.B.G. sampled the remains and B.H. did the wet laboratory work. D.G. supervised the research and wrote the paper.

## Data availability statement

All the obtained sequences are available in BAM format under bioproject PRJEB61858.

## Benefit-sharing statement

The authors declare that they had requested and got permission for the destructive bioarchaeological analyses of the archaeological material in the study from the stakeholders, excavator and processor archaeologists. All data have been shared with the broader public via appropriate biological databases.

